# Reconstructing cell type evolution across species through cell phylogenies of single-cell RNAseq data

**DOI:** 10.1101/2023.05.18.541372

**Authors:** Jasmine L. Mah, Casey W. Dunn

## Abstract

The origin and evolution of cell types has emerged as a key topic in evolutionary biology. Driven by rapidly accumulating single-cell datasets, recent attempts to infer cell type evolution have largely been limited to pairwise comparisons because we lack approaches to build cell phylogenies using model-based approaches. Here we approach the challenges of applying explicit phylogenetic methods to single-cell data by using principal components as phylogenetic characters. We infer a cell phylogeny from a large, comparative single-cell data set of eye cells from five distantly-related mammals. Robust cell type clades enable us to provide a phylogenetic, rather than phenetic, definition of cell type, allowing us to forgo marker genes and phylogenetically classify cells by topology. We further observe evolutionary relationships between diverse vessel endothelia and identify the myelinating and non-myelinating Schwann cells as sister cell types. Finally, we examine principal component loadings and describe the gene expression dynamics underlying the function and identity of cell type clades that have been conserved across the five species. A cell phylogeny provides a rigorous framework towards investigating the evolutionary history of cells and will be critical to interpret comparative single-cell datasets that aim to ask fundamental evolutionary questions.

## Introduction

Trees are a prevalent pattern in cell biology. Three distinct types of cell trees are often discussed. These are the cell lineage^1–3^, the pattern of phenotypic similarity between cells^4–6^, and the cell phylogeny^4, 7–16^. Often, these trees are conflated because the implicit null expectation is congruence. Yet the most compelling evolutionary and developmental cell biology can defy null expectations and lead to incongruent topologies^17, 18^. To explore this exciting biology, we need approaches to independently describe each of these distinct types of topologies.

Cell phylogenies are the least well-explored of the three trees. Cells are evolutionary units and homologous cell types can be readily identified across organisms^12^. A cell phylogeny depicts the evolutionary history of cells: extant cells are placed at tip nodes, and interior nodes represent cells present in ancestral organisms. Much like in a gene tree, new cell types emerge by divergences that may occur between cell types or due to speciation^15^. This contrasts with a developmental lineage, where tip nodes are differentiated adult cells and interior nodes are precursor cells that divided during embryogenesis. Branching in a lineage depicts cell division.

Cell lineages are described by well-established methods for tracking cell division^1, 3, 19, 20^ and have been extensively studied in the context of embryology^1–3^ and cancer^21–23^. Phenotypic similarity trees are likewise well-studied, as embodied in phenetic Linnaean-like cell taxonomies^4–6, 24^. However, while adjacent methods have been pursued^3, 8, 25–27^, there is far less work on building cell phylogenies using explicit evolutionary models^7, 8, 10^. Neighbour joining^28^, a hierarchical clustering algorithm, has been used to construct trees from cell phenotype data^9, 22, 29–31^. These trees describe distances between phenotypes and have been used as estimates of evolutionary history in the absence of other approaches. However, while evolutionary conservation may lead to phenotypic similarity between closely related cell types, neighbour joining trees do not explicitly model evolution.

There are multiple motivations for developing robust approaches to constructing cell phylogenies. Single-cell studies pursuing evolutionary questions across species have gained prominence, but without approaches to build a cell phylogeny most employ pairwise comparisons instead ^32–36^. Pairwise comparisons, however, can be misleading when phylogenetic structure is present^37–39^. A cell phylogeny provides a concrete framework to study fundamental evolutionary questions, such as the origins of multicellularity^40^, and will also establish clear biological criteria for defining cell types^4, 11, 12, 15^, just as phylogenetics has transformed the taxonomic organization of species. Importantly, a cell phylogeny allows us to directly apply phylogenetic comparative tools to long-held questions in cell biology^9, 27^. We can, for instance, investigate the homology between the choanoflagellate and choanocyte collars^41, 42^ with an ancestral character state reconstruction.

Here we provide an approach to building cell phylogenies that applies an explicit evolutionary model to single-cell RNAseq data from multiple species. We use principal components as phylogenetic characters and observe the phylogenetic signal in earlier components. We identify robust clades that allow us to assign a phylogenetic definition of cell type to a dataset of differentiated eye cells. Principal component loadings outline gene expression dynamics that describe the function and identity of cell type clades in the cell phylogeny.

## Methods

Several technical challenges exist in analyzing gene expression data across species. These include normalization and integration of counts across species^43–46^, the highly correlated nature of gene expression, the prevalence of noise, and the extreme multi-dimensionality of the number of characters. We approach these problems by using established single-cell RNAseq (scRNAseq) bioinformatics workflows to normalize and integrate counts^43–45^, and PCA to rotate the data and reduce rank.

Code for all analyses is available at https://github.com/dunnlab/cellphylo/.

### Data

We used single scRNAseq counts from van Zyl et al.^33^ as our pilot dataset. van Zyl et al.^33^ sampled cells of the aqueous humor of the anterior segment of the eye from five adult model species: *Macaca mulatta, Macaca fascicularis*, *Mus musculus*, *Sus scrofa* and *Homo sapiens*. This dataset was chosen for the uniformity of sampling, consistency of lab and sequencing protocols, the high quality of its cell type annotations, and the abundance of genomic resources available for the five model species. UMI counts were downloaded as CSV files from the NCBI GEO database (GSE146188). A file containing meta-data, including cluster assignment and cell type labels, was obtained from the Broad Institute Single Cell Portal^47^ (https://singlecell.broadinstitute.org/single_cell/study/SCP780/).

van Zyl et al.^33^ identified cell types by clustering their human dataset and assigning a cell type label to each human cluster according to marker gene expression and histology. They then clustered the other non-human datasets and used a machine learning algorithm to identify clusters from non-human species corresponding to the labeled human clusters. Because correspondence between clusters was not always 1:1, some non-human species had multiple clusters identified to the same human cell type cluster. Similarly, some human cell type clusters were absent in other species, while other non-human clusters were not present in humans. The latter were given their own unique cell type label. In our analyses, we considered each labeled cluster in any one species a “cell type group”, following the labeling scheme of van Zyl et al.^33^. The number of cell type groups possessed by each species ranged from 15-20.

### Normalization and integration within and across species

Standard Seurat (version 4.1.0)^45^ workflows were leveraged for normalization and integration^43–45^. For within-species analysis, SCTransform^44^ in Seurat was used to normalize the counts of each species’ matrix for cell sequencing depth and variance stabilization, using a negative binomial model of counts (Fig. S1, step 2). The Seurat function IntegrateData^43^ was subsequently used to make counts comparable across within-species sample batches (Fig. S1, step 3). The integrated Pearson residuals (from the `integrated assay`, `scale.datà slot) resulting from normalization and integration were considered in all further analyses. The total number of cells sampled per species ranged from 24,023 (human) to 5,067 (mouse), resulting in large matrices (Fig. S1, step 1). To reduce computational burden, matrices were subsetted such that each cell type group had an equal number of replicates (Fig. S1, step 4). The cell type group with the fewest replicates across all five species were mouse B cells, for which 10 cells were sequenced. Thus, when subsetting we randomly selected 10 replicate cells per cell type group in all five matrices (Fig. S1, step 4). The number of cell type groups per species ranged from 15 to 20, totalling 92 cell type groups represented by 920 cells across the five species (Fig. S1, step 4).

For cross-species analysis, Ensembl BioMart (version 107)^48^, which identifies homologs through gene trees, was used to obtain a mapping of all homologs shared between the five model species. The five matrices were then joined using one-to-one orthologs common to all five species to create a combined matrix with 920 cells and 5,870 genes (Fig. S1, step 5). One cell was removed after filtering for cells that expressed at least 200 genes and genes that were expressed in at least 3 or more cells. IntegrateData^43^ was used a second time to integrate values across species in this combined matrix, and the integrated Pearson residuals for the 2000 most highly variable genes were used for subsequent analysis (Fig. S1, step 6). Successful integration was confirmed by producing a UMAP and annotating cells by species identity and cell type identity (Fig. S2). The final matrix before rotation and rank reduction had 919 cells and 2000 genes (Fig. S1, step 6).

### Rotation and rank reduction of the data

Gene expression data is highly noisy, highly correlated, and extreme in its multidimensionality. Most models of continuous character evolution are based on Brownian motion^49^. While Brownian motion models can generate correlated data, there are multiple technical challenges to the inference of continuous data with correlation^50^. In addition, the number of parameters required to be estimated for models of correlated continuous character evolution increases quadratically with each additional character, as the mean, variance and covariances of each character must each be estimated. This results in a small N:p ratio (where N represents the number of samples and p the number of parameters), which makes parameter estimation difficult. The number of samples in our single-cell data set (919 cells, Fig. S1, step 6) are far outweighed by the number of parameters that must be calculated when data are correlated. The extensive correlation among gene expression suggests that the data may be low rank, and together with the above challenges it is therefore desirable to transform the data to this lower rank space with less correlation.

We performed a principal component analysis (PCA) on the combined cross-species integrated matrix (Fig. S1, step 6), centering the data around the mean and normalizing the variance by dividing the resulting principal components (PCs) by their standard deviation. This produced a character matrix with 919 cells and 919 PCs (Fig. S1, step 7). The rotation of our data to these new axes produced PCs that are independent and uncorrelated. We use these principal components as phylogenetic characters moving forward.

We achieved rank reduction by subsequently discarding a subset of the later PCs. Principal components are sorted in descending order by the variance they explain. Earlier principal components encode broad, slow-changing patterns of variance (often referred to as low frequency signal), while later principal components encode rapid, fine-grained variance (i.e. high frequency signal). In a phylogenetic framework, low frequency changes occur slowly across the whole topology of the phylogeny, while high frequency changes occur rapidly across all tips regardless of the greater structure of the tree. Here, we hypothesize that early principal components are enriched for phylogenetic signal, while later principal components represent cell-specific noise.

We plotted the variance described by each principal component and confirmed that the data, although high dimensional, was low rank (Fig. 1). We identified the number of principal components to retain by plotting the variance described by each principal component and identifying the elbow in the variance plot where including further principal components only minimally increases the amount of variance described (Fig. 1). This elbow occurred at PC 20, and so we retained the first 20 PCs for use as phylogenetic characters, trimming the rest.

**Figure 1.**
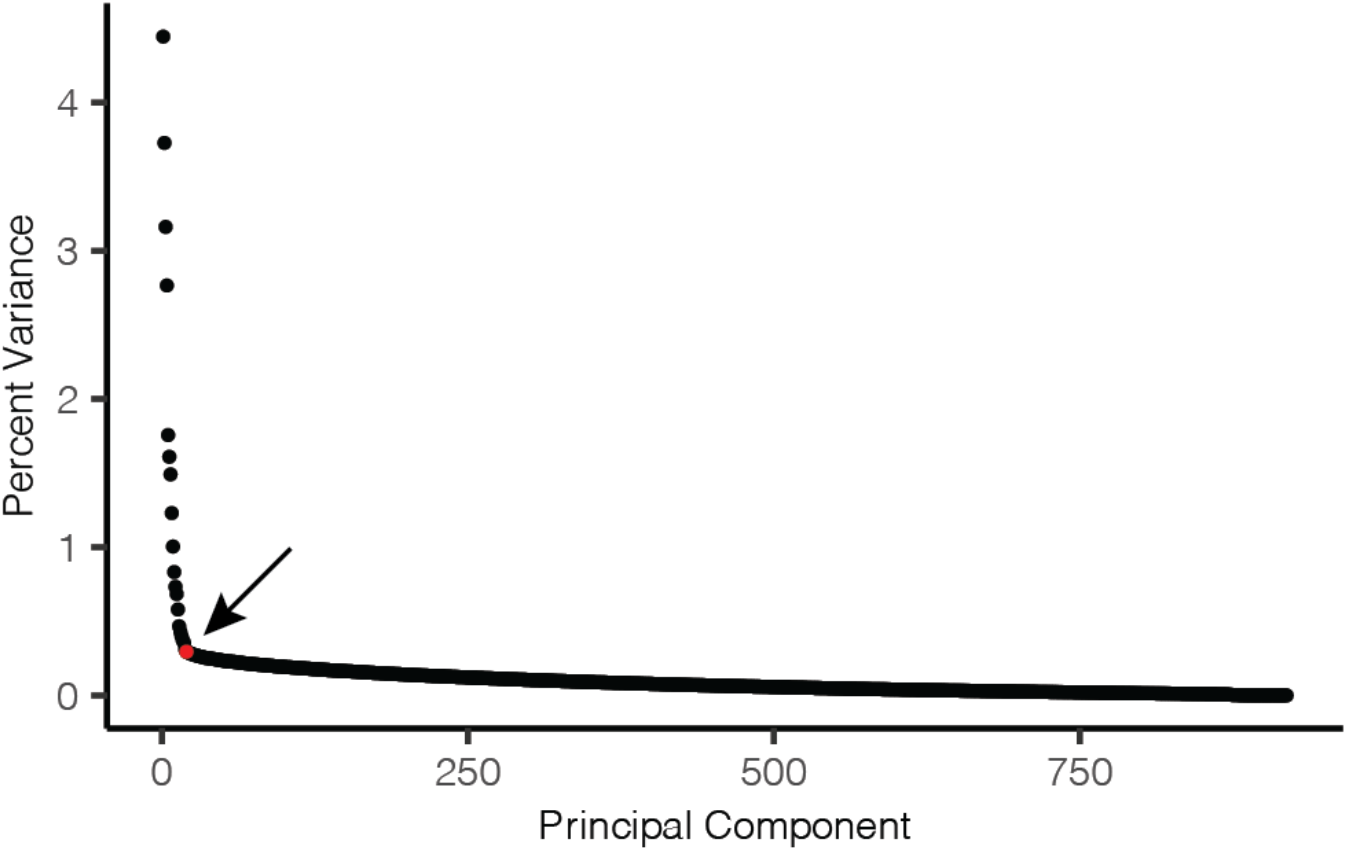
Percentage of variance described per principal component. 919 principal components were created by principal component analysis of a 919 cell 2000 gene matrix. Approximately 27% of the variance resides in the first 20 principal components, which forms a distinct elbow in the plot (arrow). PC 20 is highlighted in red.

By taking the first 20 PCs, we aim to reduce noise, enrich for phylogenetic signal and drastically reduce the number of characters from 2000 to 20. Because principal components are orthogonal and uncorrelated to each other, this allowed us to fit the data better to the uncorrelated Brownian motion model^50^ we used to infer the cell phylogeny. Our procedure also greatly simplified parameter estimation, making the model computationally feasible. Centering the data before performing PCA collapsed all means to zero, and, similarly, the use of orthogonal principal components set all covariances to zero.

### Phylogenetic inference using an uncorrelated Brownian motion model of evolution

We used an explicit model of evolution based on Browian motion to infer our cell phylogenies^49, 50^. While the use of Brownian motion-based models are well-established in the inference of phylogenies from continuous characters, like morphology ^49–53^, evolutionary models of gene expression remain underdeveloped ^54^. Phylogenetic models were originally designed upon assumptions based on mechanisms of species evolution rather than cells, and some studies indicate that the use of more complex Ornstein-Uhlenbeck (OU) models may better fit the heterogeneity of most gene expression datasets^54^. Our study is unique in that instead of using gene expression values directly, we use principal components calculated from gene expression values as our phylogenetic characters. In addition, we remove later principal components that may represent highly heterogeneous cell-specific signal.

To infer the cell phylogeny, we used contml from PHYLIP^55^ (v. 3.698, https://evolution.genetics.washington.edu/phylip.html). This program performs phylogenetic inference of continuous data by maximum likelihood search, calculating likelihoods using the REML PIC algorithm^50^, an evolutionary model explicitly based on Brownian motion.

### Principal component sweep experiment

We performed a sweep experiment to better understand the effect of the number of retained principal components on characteristics of the phylogeny (Fig. 2). To do so, we created ninety-eight 919 cell matrices that varied by the number of principal components (Fig. S1). From a 919 cell 919 PC matrix (Fig. S1, step 7), the 98 character matrices were made by serially increasing the number of PCs retained, from 3 PCs (PCs 1-3 retained) to 100 PCs (PCs 1-100 retained) (Fig. S1, step A1). It was not possible to infer a phylogeny from a character matrix that featured fewer than 3 PCs. We inferred cell phylogenies with contml using the ‘C’ (continuous character) option and leaving all other settings at default. For each of these 98 trees, we calculated total tree length (sum of all edge lengths), the sums of the tip and interior edge lengths, and the star-ness score (the ratio of the sum of tip edge lengths to the sum of interior edge lengths). These values were plotted in R^56^ (version 4.2.2) to produce Figure 2.

**Figure 2.**
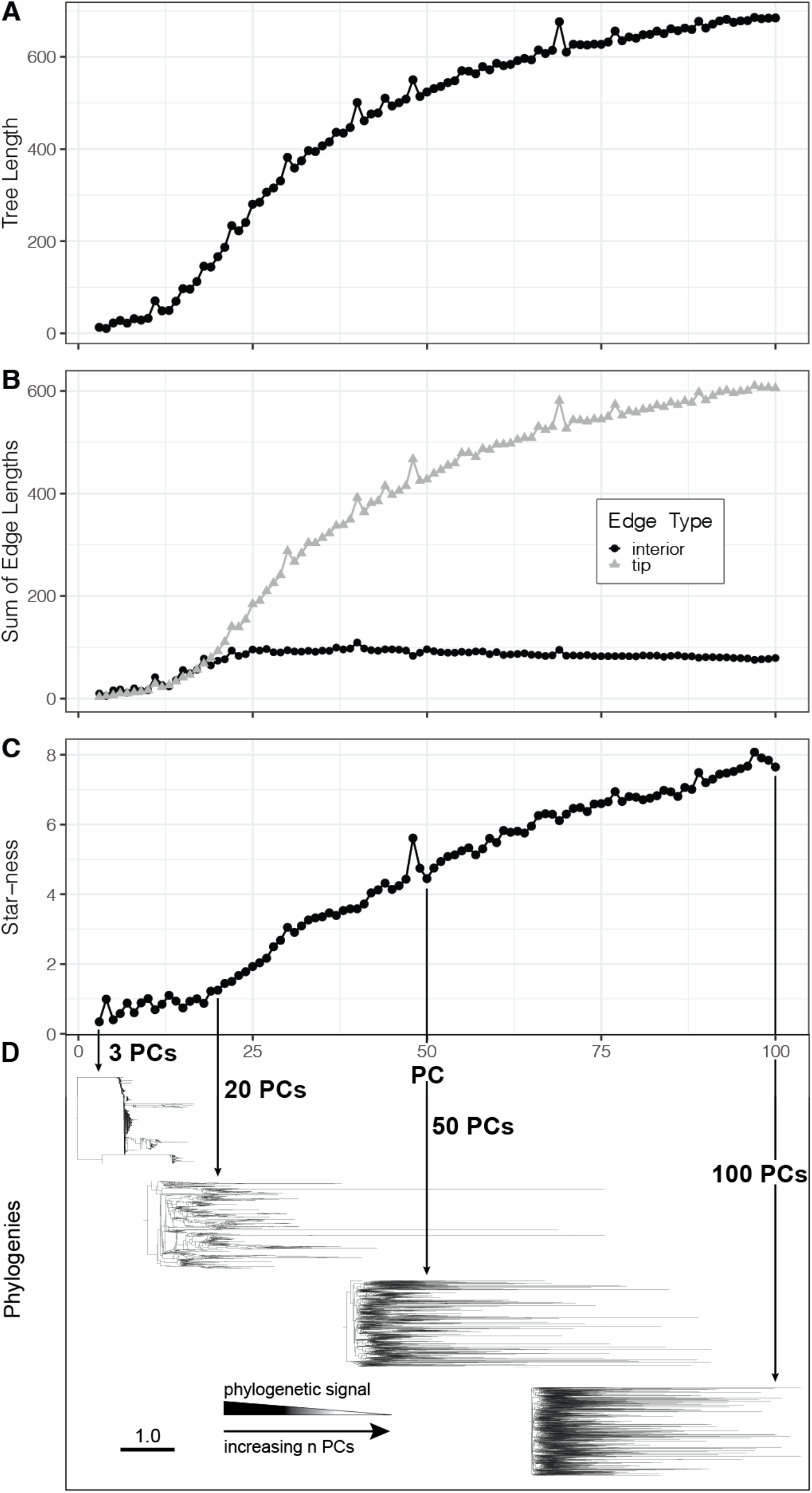
The inclusion of more principal components in the character matrix leads to a prevalence of noise, obscuring the phylogenetic signal. As the number of principal components increases, A. tree length increases, B. tip edge length increases while interior edge lengths remain constant, and C. the tree becomes more star-like. D. Examples of trees inferred from 3, 20, 50 and 100 PCs. The scale bar in D represents units of evolutionary change.

To inspect the emergence of cell type clades as the number of PCs were manipulated, we followed the above procedures and inferred a tree from a 919 cell 919 PC matrix (Fig. S1, step 7) subsetted to 20, 500 and 919 PCs (Fig. 3, Fig. S1, steps B1-3).

**Figure 3.**
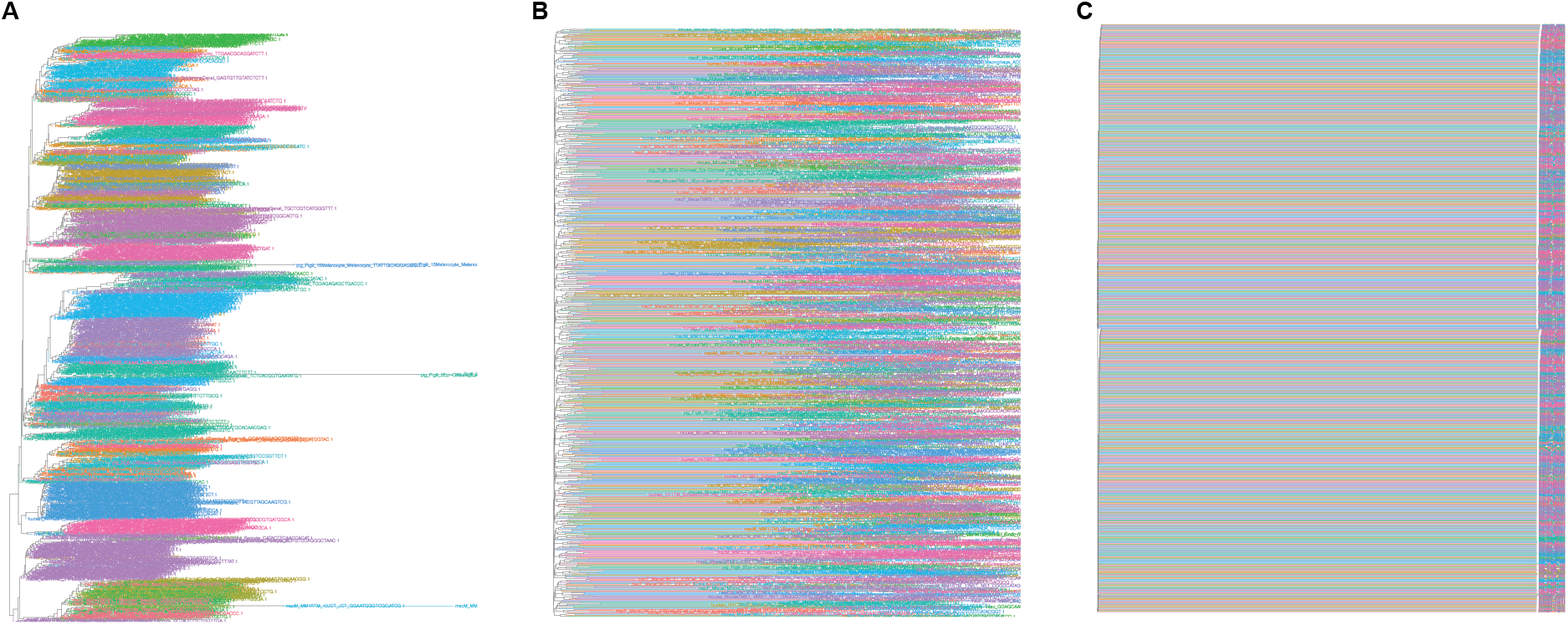
The formation of cell type clades improves when the number of principal components are reduced. Cell phylogenies of 919 cells inferred from A. 20 principal components, B. 500 principal components, and C. all 919 principal components. All three trees were inferred using the same data and parameters. Tip branches and labels are colored according to cell type. Cell type colors are consistent across all three trees.

### Inference of the focal cell-level phylogeny

We used contml to infer the focal phylogeny of Figure 4. The ‘C’ option was used to specify continuous characters were to be analyzed. The ‘G’ option was selected to search tree space by global rearrangement. Finally, the ‘J’ option (‘Jumble’) was selected to randomize the input order of the taxa 100 times and select the highest likelihood tree (see ‘*Measuring technical repeatability with jumble scores*’ for detailed discussion of the Jumble setting). All other contml settings were left at default. Finally, the cell phylogeny was midpoint rooted.

**Figure 4.**
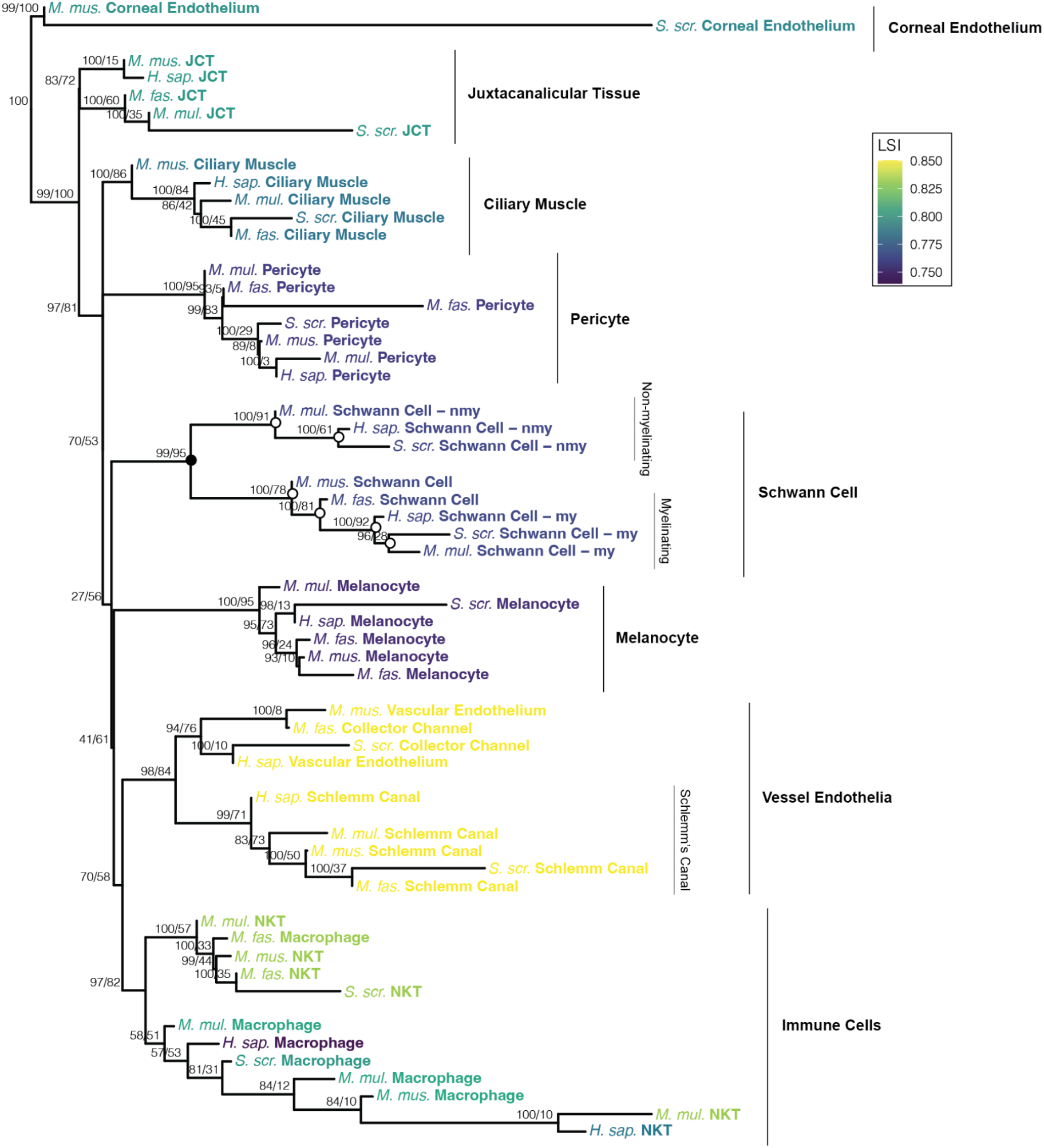
A cell phylogeny of aqueous humor cells. Fifty-four cells from five model species cluster by cell type in the cell phylogeny. Species and cell types are labeled at the tips. Cell type clades are indicated by vertical bars. Superclades include the Schwann cell, immune cells and vessel endothelia clades. In the Schwann cell superclade, the black node is an example of a within cell-type divergence, and the white nodes are examples of species-level divergences. Node support values are printed as “jumble score / scjackknife score”. The leaf stability index is plotted as tip color. H. sap., *Homo sapiens*, M. fas., *Macaca fascicularis*, M. mul., *Macaca mulatta*, M. mus., *Mus musculus*, Sus. scr., *Sus scrofa*, my, myelinating, nmy, non-myelinating, JCT, juxtacanalicular tissue, NKT, natural killer T cell, LSI, leaf stability index.

Preliminary phylogenetic inference runs were performed on a 92 cell 20 PC matrix (Fig. S1, step C1.1). We started with a 919 cell 919 PC matrix (Fig. S1, step 7). We then subsetted the principal components to 20 PCs and randomly subsetted cells to one cell per cell type group per species to create a 92 cell 20 PC matrix representing 92 cell type groups (Fig. S1, step C1.1). After initial inference on this 92 cell matrix (Fig. S4), we removed a minority of cell type groups that consistently failed to form monophyletic groups and destabilized the topology. This is similar to removing rogue taxa from a species phylogeny. These unstable cells were the corneal epithelium, beam A, beam X, fibroblast and B cells. van Zyl et al.^33^ had found that some clusters in non-human species expressed a mix of both fibroblast and beam cell marker genes, making identification of these cell types difficult to disentangle across species. Similarly, in a subset of human samples, B cells were poorly sampled and in some cases had to be identified histologically when identification by gene expression alone failed^33^. We additionally removed cell type groups represented by only a single tip, as it is difficult to meaningfully interpret the significance of such singletons in the context of other cell type clades. This produced our final 54 cell 20 PC matrix (Fig. S1, step C1.2), representing 54 cell type groups. Phylogenetic inference was performed on this matrix to produce the focal cell-level phylogeny in Figure 4.

### Measuring technical repeatability with jumble scores

The bootstrap procedure, which is commonly used to assess the repeatability of maximum likelihood trees, is built upon an assumption that each phylogenetic character is weighted equally^57^. This is not true of our characters, as principal components are weighted in descending order by the amount of variance described. This means bootstrapping is not appropriate for our application.

Instead, we assessed the repeatability of the topology by perturbing the starting position of the maximum likelihood search in tree space. Should phylogenetic inference be repeatable, the same topology should be inferred regardless of where the maximum likelihood search was initiated. This perturbation was achieved by using the ‘J’ option (‘Jumble’) of contml, which created different starting trees by randomizing the input order of the taxa. The contml options we used are ‘C’ (continuous characters), ‘J’ (jumble 100x), ‘G’ (global rearrangement), and all other settings at default. We performed 250 contml runs on the same dataset, jumbling 100 times each run and inferring a phylogeny from each jumble. This produced 100 ‘jumble trees’ per run, from which the phylogeny with the highest likelihood was selected and presented as the final output of the run. A percentage (5.2%) of contml runs failed, leaving us with 237 high likelihood jumble trees. We then calculated the clade frequencies of these 237 jumble trees using plotBS (with type = ‘phylogram’ and method = ‘FBP’) from the phangorn^58^ R package (version 2.10.0). These frequencies were plotted onto the nodes of the focal cell-level phylogeny of Figure 4 as ‘jumble scores’, which represent the percentage of jumble trees in which that split was present.

The focal cell-level phylogeny was selected from among the 237 jumble trees by identifying the phylogeny with the greatest overall sum of jumble scores. Fourteen phylogenies fit this criteria. They possessed identical topologies (Robinson-Foulds distance^59^ = 0) or minimal differences when branch length was taken into account (average branch-score-distance^60^ = 0.0298). The tree with the smallest sum of branch-score-distances between the fourteen trees was selected as the focal phylogeny presented in Figure 4.

### Measuring biological repeatability with single-cell jackknife scores

We expect that cell gene expression may exhibit significant biological variability, even among cells of the same cell type group, and thus needed to measure the robustness of the topology to cell sampling. We assessed biological repeatability using a single-cell jackknife (‘scjackknife’) procedure. This was made possible by the fact that single-cell data sets provide multiple replicate cells per cell type group. We performed a jackknife procedure, where we randomly drew 1 cell per cell type group per species from a 919 cell 20 PC matrix (Fig. S1, step C2.1), which possesses 10 cells per cell type group. We selected cells from among the 54 cell type groups that were used to infer the phylogeny in Figure 4. This produced a jackknifed matrix with 54 resampled cells and 20 PCs (Fig. S1, step C2.2). We performed this jackknife procedure 500 times to create 500 new 54 cell matrices (Fig. S1, step C2.2), where the specific identity of the cell for a particular cell type group had been randomly sampled. The number of PCs were kept consistent at 20 PCs. From each matrix, a phylogeny was inferred with contml using the same settings as described above for the jumble trees. After removing runs that errored, we retained 439 high likelihood scjackknife trees.

We used the transfer bootstrap expectation (TBE) score^61^ to measure the consistency of the topology across the scjackknife trees, and used this as a measure of support for biological repeatability. The TBE is similar to Felsenstein’s bootstrap^57^ where the frequency of a clade is calculated across a set of trees, except instead of scoring the presence of a clade through a simple presence/absence index, it uses transfer distance to identify clades that are present but exhibit some variability across comparison trees. Transfer distance between a branch in the reference tree and a branch in the comparison tree is equal to the number of branches that must be removed (‘transferred’) to make the two branches equivalent^61^. This is useful for data sets that exhibit clear topological patterns, but whose clades exhibit small variation not captured by a boolean presence/absence procedure. In Felsenstein’s bootstrap, even if all but one taxon is present in the comparison clade, the bootstrap score is 0, which is not reflective of the high degree of similarity shared by the two clades (Fig. S5). We do not expect that the stability exhibited by species trees will be matched by a cell phylogeny. The cell phylogeny possesses taxa that exist at an entirely different level of organization and is built using highly variable gene expression, rather than molecular sequence data. Likewise, there is no precedence for choosing a significance threshold in a cell phylogeny. Lemoine et al.^61^ found that a TBE threshold of 70% reliably identified branches concordant with the true tree of a species phylogeny built from molecular sequence data, and so we use this threshold to conservatively define well-supported clades. We calculated the TBE from the 439 scjackknife trees using Booster (https://github.com/evolbioinfo/booster) and mapped these scores onto the focal phylogeny in Figure 4 as ‘scjackknife scores’.

### Leaf stability index

The leaf stability index^62^ was calculated from the set of jumble trees produced for Figure 4 using phyutility^63^, with the command: ‘java -jar phyutility.jar -ls -in jumble_trees.tre’. These values were plotted by colour onto the cell phylogeny in Figure 4.

### Creating a cell phylogeny from averaged cell type group replicates

The 919 cell 20 PC matrix (Fig. S1, step C2.1) has ∼10 replicate cells per cell type group (92 cell type groups). We followed the above procedure to infer a tree and calculate jumble and scjackknife scores, but instead of selecting a single representative cell per cell type group we averaged values across multiple replicates. For the jumble trees, we averaged the PC loadings of all ten replicates, selecting the 54 cell type groups used in Figure 4. This produced a matrix with 54 averaged cell type groups and 20 PCs (Fig. S1, step D1). We performed 500 jumble tree runs and received 209 high likelihood jumble trees, where each tip is a cell type group represented by the average of 10 replicate cells. For the scjackknife trees, we randomly sampled 5 cells for each cell type group from the 919 cell 20 PC matrix (Fig. S1, step C2.1), selecting for the 54 cell type groups used in Figure 4. Their PC loadings were averaged to create a new jackknifed matrix, with 54 averaged cell type groups and 20 PCs (Fig. S1, step D2). We created 500 scjackknife matrices and received 393 high likelihood scjackknife trees, where each tip indicates a cell type group represented by the average of 5 randomly chosen replicates. We used the best scoring jumble tree to plot jumble and scjackknife scores onto in Figure 5. Five jumble trees tied for the highest sum of jumble scores; the most topologically representative among this subset was selected as the “best” jumble tree following the same procedure as described for Figure 4.

**Figure 5.**
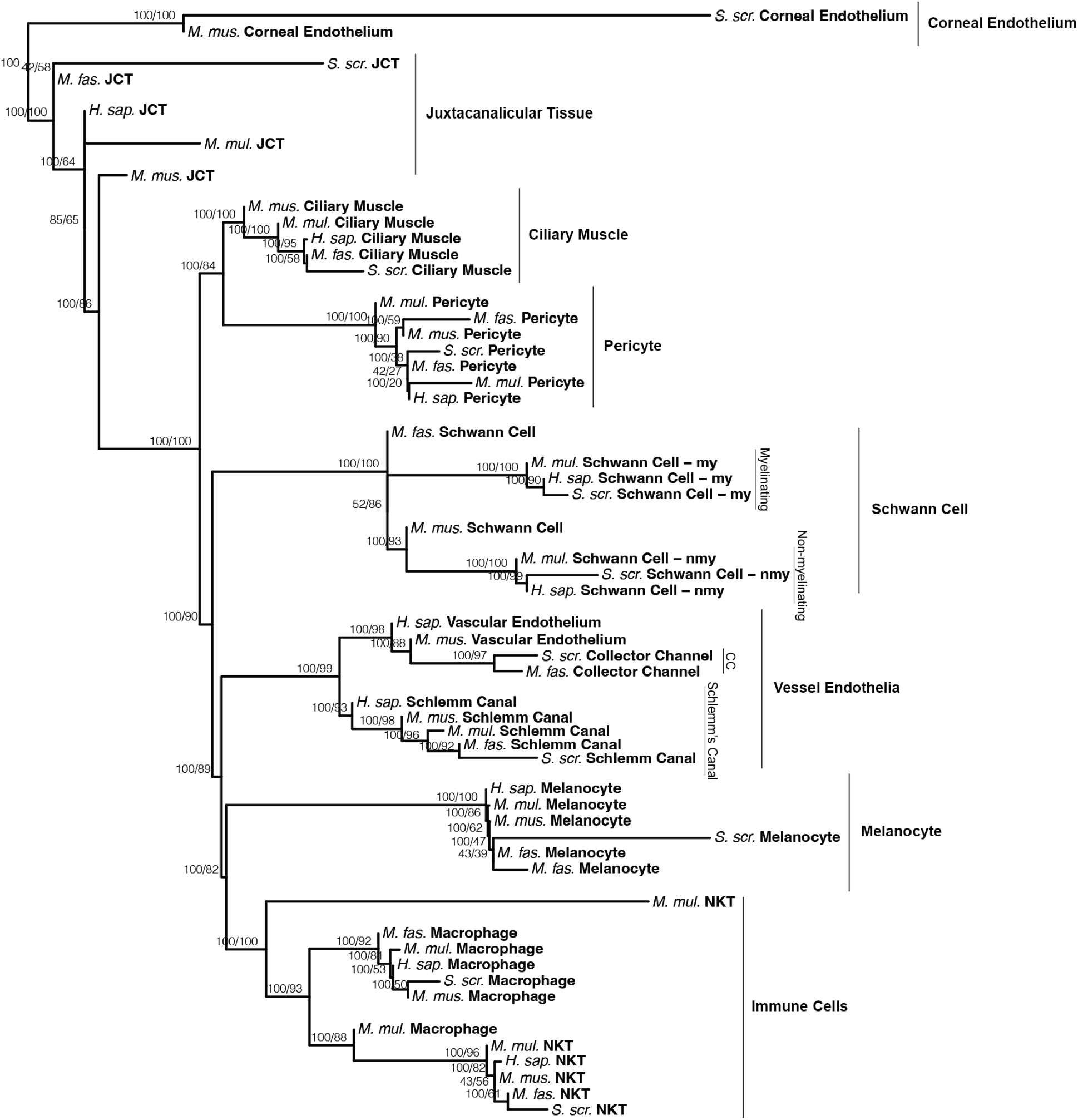
Averaging replicate cells resolves relationships between and within cell type clades. In this phylogeny, each tip corresponds to a cell type group represented by the average of ten cells. High jumble and scjackknife scores (plotted as “jumble score/scjackknife score”) characterize many nodes, including those along the backbone of the cell phylogeny. Cell type clades present in Figure 4 are also present here and indicated with vertical bars. Species and cell types are labeled at the tips. H. sap., *Homo sapiens*, M. fas., *Macaca fascicularis*, M. mul., *Macaca mulatta*, M. mus., *Mus musculus*, Sus. scr., *Sus scrofa*, my, myelinating, nmy, non-myelinating, JCT, juxtacanalicular tissue, CC, collector channel, NKT, natural killer T cell

### Identifying genes with high principal component loadings

We examined the identity of the genes with the highest positive and negative loadings for the first 20 PCs from the results of the PCA that produced the 919 cell 919 PC matrix (Fig. S1, step 7). We extracted the gene loadings, identified genes by their human gene symbol and sorted them by most positive and most negative loading scores for each PC. For the top 10 most positive or most negatively loaded genes for each PC, we manually investigated known functions of the gene using the NCBI gene database and/or a literature search. These genes and descriptions are summarized in Supplementary File S1 for positive loadings and Supplementary File S2 for negative loadings. A summary of the cell type signals observed amongst the gene loadings is provided in Table 1 and Supplementary File S3. The unscaled 919 cell 919 PC matrix is provided as Supplementary File S4.

**Table 1.** Cell type-specific signals are present among the most highly loaded genes in early principal components. Gene loadings for the first ten principal components were ordered by most positive or negative loading. The top ten most positive or negative loadings were enriched for genes specific to the development, function and identity of the specific cell type clades that emerged in the cell phylogeny. The correspondence between PC loading and cell type clade signal is provided in this table. Loadings that had no immediately apparent cell type affinity are labeled ‘?’.

| Principal Component | Positive Loading | Negative Loading |
| --- | --- | --- |
| PC1 | immune | muscle |
| PC2 | vessel endothelium | neural system |
| PC3 | ? | muscle |
| PC4 | fibroblast | melanocyte |
| PC5 | neural system | immune |
| PC6 | ? | ? |
| PC7 | ? | neural system |
| PC8 | ? | ? |
| PC9 | immune - myeloid | immune - lymphoid |
| PC10 | vessel endothelium | ? |

## Results

### Single-cell gene expression data is low rank, with first principal components containing phylogenetic signal

The combined and cross-species integrated matrix consisted of 919 cells and 2000 genes (Fig. S1, step 6). This matrix represented 92 cell type groups across five species, each with 15-20 cell type groups, for which ∼10 cells were sampled per cell type group. Following integration, a UMAP revealed that cells clustered by cell type rather than species, consistent with successful cross-species integration (Fig. S2).

Gene expression is noisy, correlated and highly multidimensional. To apply an uncorrelated evolutionary model for phylogenetic inference, we performed a PCA and used the principal components (PCs) of gene expression as phylogenetic characters. PCs are orthogonal, and both dimensionality and noise can be reduced by trimming later PCs if the data is low rank. Plotting the amount of variance described per principal component revealed a clear elbow at PC 20 (Fig. 1). Approximately 27% of the variance is explained by the first 20 PCs, while there was negligible change in the amount of variance explained by the remaining 899 PCs (Fig. 1). This demonstrates the data is low rank.

We used a Browian motion model of continuous character evolution^50^ to infer cell phylogenies from PCs derived from gene expression levels. We inferred unrooted phylogenies, where each tip is a cell type group represented by a single cell and branch lengths indicate the amount of evolutionary change in expression. We examined the impact of rank on the cell phylogeny by inferring trees from matrices with an increasing number of PCs (Fig. 2). Trends in tree length, tip v.s. internal edge length and star-ness of the phylogenies were observed. The plots of these trends exhibited elbows after PC 20 (Fig. 2), corresponding to the elbow in the principal component variance plot (Fig. 1).

Total variance can be measured as tree length. Since PCs are sorted in descending order of variance (Fig. 1), tree length increased most rapidly as early PCs were added (PCs 3-25), while growth in tree length slowed as later PCs were included (Fig. 2A).

We characterize the added variance by inspecting the behavior of tip edge lengths and interior edge lengths as the number of PCs was increased (Fig. 2B). Phylogenetic signal, the variance that is shared among tips due to common ancestry, is represented as interior edge lengths in a phylogeny. In contrast, cell-specific variance, which may include cell-specific evolution, cell cycle state, and observational noise, is reflected in tip edge length. We hypothesized that much of this cell-specific variance is noise, and if so, is present as the higher frequency signal that resides in later PCs. We found that tip edge length increased while interior edge lengths remained relatively constant as the number of PCs increased (Fig. 2B), suggesting that later PCs describe high frequency cell-specific signal.

Finally, we examined how adding higher frequency signal affected the phylogenetic signal of the phylogeny (Fig. 2C). Star phylogenies, which have long tip edges and negligible interior edge lengths, possess little to no phylogenetic signal. We find that by adding later PCs, the phylogeny became more star-like (Fig. 2C). This suggests that adding higher frequency variance from later PCs drowns out the phylogenetic signal.

Clustering of cells into cell type clades improved significantly when the number of PCs used to infer the phylogeny was reduced (Fig. 3). Clear grouping by cell type clade could be seen in the tree made from 20 PCs (Fig. 3A). A phylogeny inferred from 500 PCs exhibited poor grouping (Fig. 3B), while one inferred from all 919 PCs failed to produce cell type clades (Fig. 3C).

### Identification of phylogenetic cell type clades

Based on the above trends, we inferred an unrooted branch-length phylogeny of 54 cells (Fig 4). This described 54 cell type groups (1 cell per cell type group) from five species, each of which possessed 15-20 cell type groups (Fig. 4). We used the first 20 PCs calculated from gene expression as phylogenetic characters (Methods). In this phylogeny, cells from across the five species clustered by cell type despite varying branch lengths, indicating the presence of a strong cell type signal that was not overwhelmed by species-level or cell-specific signal (Fig. 4). In addition, cells clustered by cell type regardless of the expression of key marker genes that have traditionally been used to identify particular cell types. For instance, the pig JCT cell, which exhibits a long branch (Fig. 4), does not express CHI3L1, a key marker gene expressed in human, macaques and mouse JCT cells^33^. Similarly, Schlemm’s canal cells from pig and mouse exhibit more dominant expression of lymphatic marker genes than in humans and macaques^33^, but still formed a single well-supported Schlemm’s canal cell clade (Fig. 4).

Just as phylogenies have directly informed species taxonomies, the cell phylogeny allows us to phylogenetically define cell type by clade membership (Fig. 4). These cell type clades included corneal endothelium, JCT, ciliary muscle, pericyte, myelinating and non-myelinating Schwann cells, melanocyte, and Schlemm’s canal cells. As indicated by the node colored with a black circle in Figure 4, myelinating and non-myelinating Schwann cells were sister to each other, forming a Schwann cell superclade (Fig. 4). While macrophages and natural killer T cells did not perfectly segregate into separate clades, they formed a single robustly-suppo rted superclade of immune cells (Fig. 4). Finally, a clade of vascular endothelia and collector channel cells were present as a sister group to Schlemm’s canal cells (Fig. 4). We label the superclade encompassing these three cell types as the ‘vessel endothelia’ clade, following van Zyl et al.^33^. Ciliary muscle cells, pericytes, and the clade encompassing Schwann cells, melanocytes, vessel endothelia, and immune cells formed a polytomy (Fig. 4).

### Robustness of the cell phylogeny

We next aimed to measure the repeatability of the topology of the cell phylogeny. Because bootstrap scores were not appropriate for our data^57^, we assessed technical repeatability with ‘jumble scores’ that summarize consistency of relationships given different addition orders of tips. Like bootstrap scores, jumble scores nearing 100% indicate that the split is highly repeatable across a set of jumble trees that were produced by repeatedly perturbing the maximum likelihood search. We found high support for splits that defined cell type clades, with most scores greater than 95% (Fig. 4). The subclades of the Schwann cell and vessel endothelia superclades were also well-supported, giving validity to their status of being closely related in a superclade relationship (Fig. 4). Several of the deeper splits along the backbone displayed much more variation across the jumble trees, leading to scores below 70% (Fig. 4). Because of this, broader relationships between clades at these splits generally remain poorly characterized, even as the cell type clades themselves are robustly supported (Fig. 4).

Given that inference was performed on a small subset of cells from the full matrix and that gene expression may vary even across cells of the same cell type, we also assessed the effect of cell subsampling on topological stability. Unlike most phylogenetic data sets we have a large pool of biological replicates, allowing us to perform a single-cell jackknife (scjackknife) procedure. The cells at the tips of the 54 cell phylogeny (Fig. 4) were repeatedly randomly resampled from a larger data set, to create 439 scjackknife trees that varied in the identity of the sampled cells. Calculating the transfer bootstrap expectation^61^ score for clades in the cell phylogeny from this set of scjackknife phylogenies revealed consistent support (more than or equal to 70) for the cell type clades that were robustly supported by jumble scores (Fig. 4).

### Tip stability is clade-specific

We assessed the topological stability of tips in the cell phylogeny by calculating the leaf stability index^62^ (LSI) from the jumble trees (Fig. 4, S3). This index ranges from 0 (very low stability) to 1 (very high stability). We found that LSI varied closely by cell type clade and was highly concordant within cell type clades, suggesting that instability is cell type-specific and describes stability of the clade as a unit (Fig. 4, S3). The most stable cell type was the Schlemm’s canal cells whereas the least stable were the melanocytes (Fig. 4, S3). The LSI values of the Schwann cell types, vascular endothelial and collector channel cells all had a standard deviation of zero (Fig. S3).

### Averaging replicate cells resolves relationships between and within cell type clades

We inferred cell phylogenies where each tip indicates a cell type group represented as the average of multiple replicate cells, rather than as a sample of a single cell (Fig. 5). Almost all jumble scores in the resulting phylogeny were 100 and most scjackknife scores fell above 70, even amongst deep nodes. The relationships within the cell type clades were also better resolved. For instance, the collector channel cells now formed a clade nested within vascular endothelia, with high jumble and scjackknife support (Fig. 5). The JCT cells were paraphyletic, but with low scjackknife support (Fig. 5). In Figure 4, the ciliary muscle and pericyte clades were in a polytomy, but here they emerged as well-supported sister clades (Fig. 5).

### Cell type signals are present among the highly loaded genes of the principal components

To identify the genes whose expression variance contributes most to phylogenetic signal, we examined the top ten most highly loaded genes of the twenty PCs that were used to infer the cell phylogeny. In several cases, genes that have been previously associated with particular cell types based on functional work and previous expression analyses have high loadings in the first PCs (Table 1). Particular cell type clades, such as the vessel endothelia clade, are reflected in the upregulation of positively-loaded clade-specific marker genes. For instance, the positive loadings for PC 2 were dominated by vasculogenesis-related genes, including validated marker genes^33^ for the Schlemm’s canal, collector channel and vascular endothelial cells (e.g. PCAM-1) (Table 1 and Table S1, Supplementary File S1). Other cell type signals were present among the negative loadings, suggesting that the phylogenetic identity of these cell type clades was defined by downregulation of their marker genes among other cells. One example is the melanocyte clade, defined by high negative loadings of melanocyte-specific genes in PC 4, including the melanocyte master transcription factor MITF^33, 64, 65^ (Table 1 and Table S2, Supplementary File S2).

## Discussion

Our findings suggest that the first principal components (PCs) of single-cell gene expression data describe evolutionary variance across cells and that this variance can be used to construct cell phylogenies. Cells from distantly-related species form well-supported cell type clades, which emerge only after removing the large number of low-variance PCs that describe fine, cell-specific differences. Liang et al.^8^ was the first to statistically demonstrate tree-like evolutionary structure among cell transcriptomes. We aimed to build from that foundation by applying an explicit phylogenetic approach using an evolutionary model. Our ability to construct a robust cell phylogeny from single-cell expression levels is consistent with Liang et al.’s^8^ finding that there is phylogenetic structure among cell transcriptomes^7–9^.

The emergence of robust cell type clades in the cell phylogeny enabled us to provide a phylogenetic definition of cell type to the cells of the tree. Marker gene expression is traditionally used to identify cell types but most marker genes have been identified and characterized in model organisms, especially humans. We find that in species distantly related to humans, phylogenetic signal is sufficient to identify such cells by membership in a cell type clade, even when these cells did not express these marker genes (e.g. pig JCT cells^33^). The topology of the cell phylogeny provides a rich biological rationale to a phylogenetic classification of cells as these relationships describe the evolutionary mechanism by which new cell type identities arise.

While there are many processes that may create branching patterns, several observations indicate that the signal in the first PCs is evolutionary variance across cells that is appropriately analyzed in a phylogenetic context. A ‘cell type’ is an identity^11, 13^ that transcends cell states, transient physiological conditions, tissue contexts, life histories, and correlated within-species evolution^46^. Despite these possible signals, here we find that robust cell type clades emerge across five distantly-related species (Fig. 4, 5). Additionally, the leaf stability index scores indicate that variance in the phylogenetic signal is uniquely specific to the cell type clade: cells of the same clade exhibit phylogenetic instability as a unit, possibly moving around the tree together as a clade (Fig. 4, S3). This provides a unique “meta-phylogenetic” character independent from phenotypic similarity that can be used to characterize cell types, further supporting the utility of phylogenetic signal to define cell types. Finally, the presence of clear cell type-specific signals among the gene loadings of the PCs from which the cell phylogeny was inferred reveals a biological logic underlying the emergence of cell type clades in the topology (Table 1, S1 and S2). This allowed us to identify the expression dynamics of genes responsible for the broad patterns of variation that defined the function and identity of conserved cell type clades (Table 1, S1, S2, Supplementary Files S1-3).

Consideration of all three trees of cell biology - the developmental lineage, phenotype similarity tree, and cell phylogeny - is essential to address the most fundamental questions about cell diversity and evolution. There is a long record of tools for the study of developmental or cancer cell lineages^1, 19, 31^ and phenotype similarity^66–71^, but cell phylogenies have seen much less rigorous attention despite rapidly growing interest ^4, 8–10, 15, 29, 72, 73^. Here we provide an approach to construct the cell phylogeny. RNAseq expression data is information-rich, but the ability to construct a cell phylogeny from expression levels is challenged by noise, covarying expression, and high dimensionality. We find that rotation and rank reduction of the data can overcome these challenges and make the data more suitable for analysis with existing continuous trait phylogenetic tools that assume character independence^49, 50^. While Brownian motion is a simple model of evolution, our use of it was able to produce the emergence of well-supported cell type clades (Fig. 4, 5), suggesting that it was sufficient for our needs.

The topology of a cell phylogeny allows us to infer specific events in the history of cell type evolution. The node at the base of the tree is the ancestral cell in a common ancestor, which then differentiated in the course of evolution to give rise to multiple distinct cells (Fig.4). Other interior nodes allow us to infer the mechanism by which new cell types emerge. Much like in a gene tree, some interior nodes represent divergences between cell types and others divergences between species (Fig. 4). The node defining the Schwann cell superclade is an example of a within-cell type divergence; an ancestral Schwann cell type program split into the myelinating and non-myelinating subtypes (Fig. 4, black node). There was subsequent divergence through speciation events, in which both Schwann cell subtypes each split with the divergence of the five species (Fig. 4, white nodes). These evolutionary events provide the biological reasoning behind a phylogenetic definition of cell types^12^. The sister clade relationship of the myelinating and non-myelinating Schwann cell types suggest that these two cell types may have arisen as sister cell types during animal evolution^11–13^.

The topology of the cell phylogeny is also consistent with hypothesized scenarios of cell type evolution^74^. Animal cell type evolution can be traced back to the emergence of the three germ layers and different germ layer origins is a fundamental distinction between cell types^75, 76^. Eye development involves a complex series of events involving ectoderm-derived neural crest cells and mesoderm-derived cell types^77^. While vessel endothelia and corneal endothelia are both described as endothelia, they are not unified in a single clade in the cell phylogeny (Fig. 4, 5). Accordingly, the corneal endothelium has a distinct embryological origin from vessel endothelia, arising not from mesoderm but from the first wave of neural crest cells during eye development^77^. The corneal endothelium is among the earliest tissues to differentiate during eye development^78^ and is placed as sister to all other cells in the rooted cell phylogeny (Fig. 4, 5).

More fine-grained evolutionary stories are also present in the cell phylogeny. Our phylogeny hypothesizes a close evolutionary relationship between the Schlemm’s canal (SC), collector channel (CC) and vascular endothelium cells, which emerged as a robustly-supported superclade (‘vessel endothelia’, Fig. 4). Some early work suggested a neural crest origin of the SC^79, 80^, but embryological studies have since traced its emergence from the limbal blood vessels^81^, congruent with its grouping with mesoderm-derived cells in the vessel endothelia clade. CCs are channels that are continuous with the SC, directly connecting the SC to the aqueous veins. Less work has been performed on their developmental origins, but their anlage appears to arise from straight vessels that sprout from the outer wall of the SC^82, 83^. It has also been hypothesized that the CCs develop from the intrascleral venous plexus ^84, 85^. Despite this close anatomical and functional relationship, the cell phylogeny hypothesizes that the CC is more closely related to the vascular endothelia of capillaries instead of the SC, with the CC forming a clade nested within the vascular endothelia (Fig. 5). Vertebrates possess two vascular systems - the blood and lymphatic systems. Mesoderm gives rise to blood vessel endothelia, which may then transition to a lymphatic fate by the expressions of specific marker genes, like PROX1 ^86^. The expression of these lymphatic marker genes in the SC (eg. PROX1, CCL21, FLT4/VEGFR3)^33^ has led to the emerging hypothesis that the SC is a unique vessel with some affinity to the lymphatic system^81, 87^. CC do not express these genes - instead they express venular markers (eg. ACKR1)^33^. The topology of the cell phylogeny is consistent with the venous v.s. lymphatic split of the two vascular systems during vertebrate evolution and development, with the CC being of venous origin, while the SC may have a lymphatic association.

When the tips of the cell phylogeny are averages of replicate cells, topological stability within and between cell type clades significantly increased (Fig. 5). The pericytes and ciliary muscle (CM) cells, both neural crest-derived, are well-supported sister clades (Fig. 5). However, in the eye they hold notably distinct biological roles^77^. While their link is understudied, extensive parallels have been drawn between pericytes and vascular smooth muscle cells (vSMCs)^88–90^. Both are contractile mural cells that regulate blood flow^88^. In some cases they differentiate from the same developmental precursor^91, 92^ and share key smooth muscle marker genes, including ACTA2^33, 88^. It is in this smooth muscle identity that significant similarities between pericytes and ciliary muscle lie. ACTA2 is also selectively expressed by CM^33^ and the canonical vSMC regulators NO^93^ and endothelin-1^94^ govern the contraction of both CM^95, 96^ and pericytes^97^. Neural crest cell types present a unique challenge due to potentially diverse evolutionary origins, but these similarities affirm a close evolutionary relationship between pericytes and CM, consistent with phylogenetic topology (Fig. 5).

All cross-species studies are implicitly evolutionary^37–39^, but despite growing excitement in comparative scRNAseq we still lack a true phylogenetic comparative approach to cell biology. Here we provide a means that will enable an explicit phylogenetic approach to evolutionary questions involving potentially any cells that can be described by scRNAseq data. Not only does this allow the leverage of phylogenetic comparative methods directly to cell biology, but the ability to infer the cell phylogeny completes the triptych of trees describing the cell lineage, phenotype similarity and evolutionary history of cells. These three trees, with their distinct but mechanistically-related topologies, have long existed in separate fields of study. However, it is their deconvolution and then systematic comparison that will give rise to insights that cannot be achieved by any of the trees alone, as the most compelling biology exists where they are incongruent. For example, one principle of development observed in *C. elegans* is that cell transcriptomes exhibit an initial “cell lineage signal” earlier in development^17^. However, as cell division progresses, this signal fades and the transcriptomes of cells of the same cell type, which are characterized by shared phenotypic characteristics, converge when adult terminal differentiation is approached^17^. This striking divergence in the topologies of the developmental lineage and phenotypic similarity trees is causally linked by the cell phylogeny: phenotypic similarity arises from shared evolutionary ancestry, not development. The cell phylogeny will enable us to deconvolute the intertwined branching patterns of phenotype and development, while providing a unifying mechanistic understanding of the emergence of cell type diversity at both the developmental and deep evolutionary time scales. Together, as the topologies of these three trees are systematically explored, a phylogenetic approach to cell biology will provide a path to connect the branching pattern of cell evolution to the broader tree of life.

## Supplementary Materials

Code and supplementary files can be found in the git repository: https://github.com/dunnlab/cellphylo

## Acknowledgements

We would like to thank members of the Dunn lab, Dr. Günter Wagner, Dr. Jacob Musser, Daniel Stadtmauer, Dr. Arun Chavan and Dr. Dean Adams for insightful comments that greatly improved our manuscript and analyses.

