## Supplementary_Figures for "Reconstructing cell type evolution across species through cell phylogenies of single-cell RNAseq data"

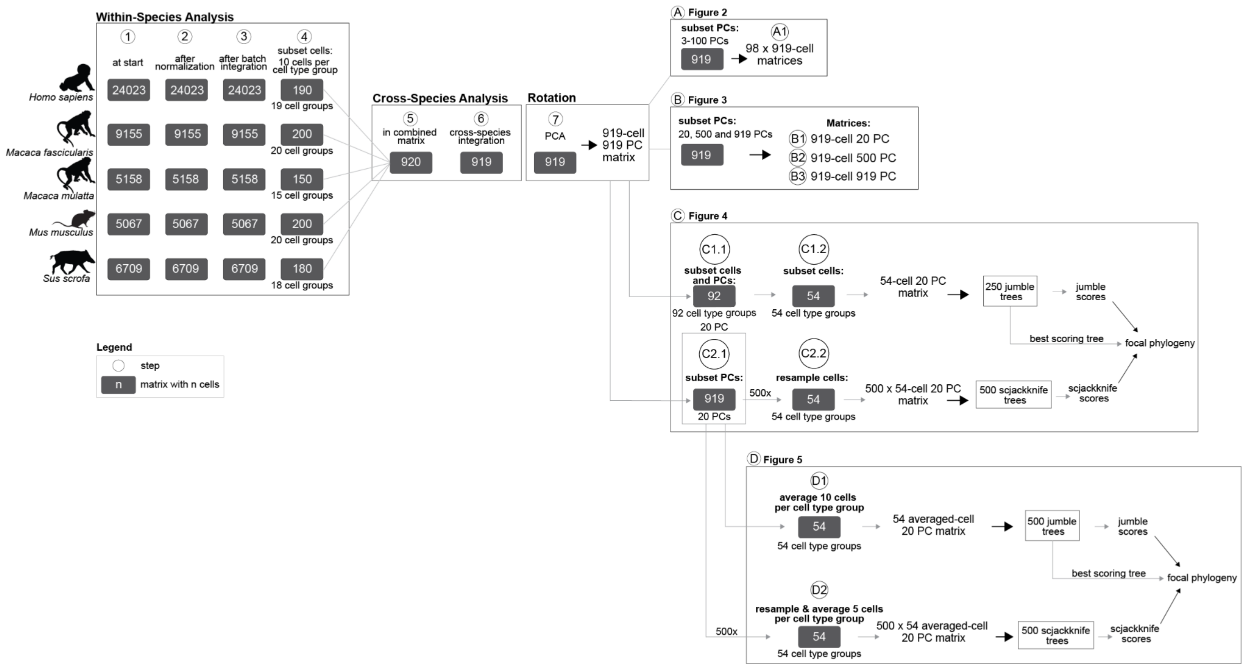


**Figure S1. Matrices created at each step of the workflow.** Each grey box represents a matrix, within which the number of cells is indicated. Matrices were first processed separately for each species ("Within-Species Analysis" - steps 1-4), then later combined into a single multi-species matrix ("Cross-Species Analysis" - steps 5-6) and rotated (step 7) to produce a 919 cell 919 principal component matrix. The matrices used for Figures 2-5 were subsequently built by subsetting from this matrix. The animal silhouettes were obtained from PhyloPic[98].

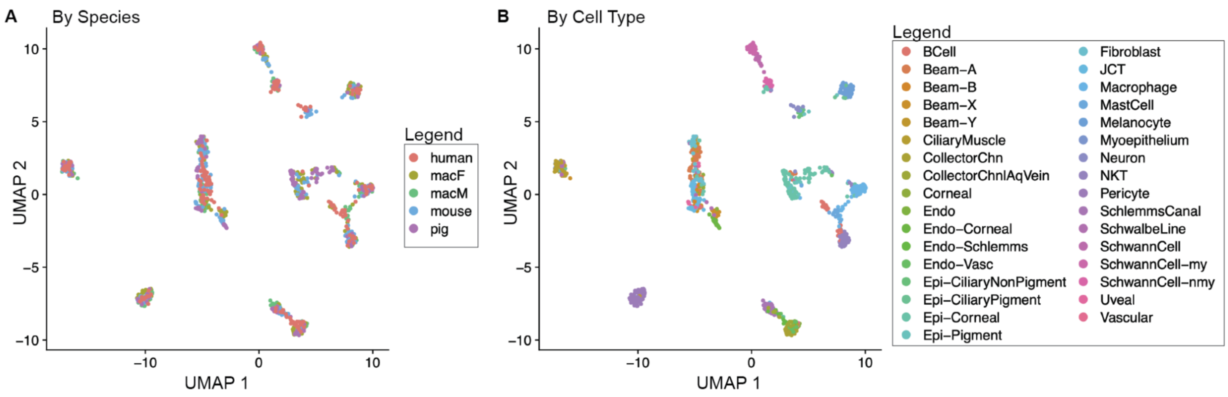


**Figure S2. Cells cluster into multi-species cell type clusters after cross-species integration**. A UMAP plot of cells, colored by A. species identity and B. cell type identity. macF, *Macaca fascicularis,* macM, *Macaca mulatta,* CollectorChn, collector channel cell, CollectorChnAqVein, collector channel aqueous vein cell, Endo, endothelium, Endo-Corneal, corneal endothelium, Endo-Schlemms, Schlemm's canal endothelium, Endo-Vasc, vascular endothelium, Epi-CiliaryNonPigment, non-pigmented ciliary epithelium, Epi-CiliaryPigment, pigmented ciliary epithelium, Epi-Corneal, corneal epithelium, Epi-Pigment, pigmented epithelium, JCT, juxtacanalicular tissue, NKT, natural killer T cell, my, myelinating, nmy, non-myelinating.

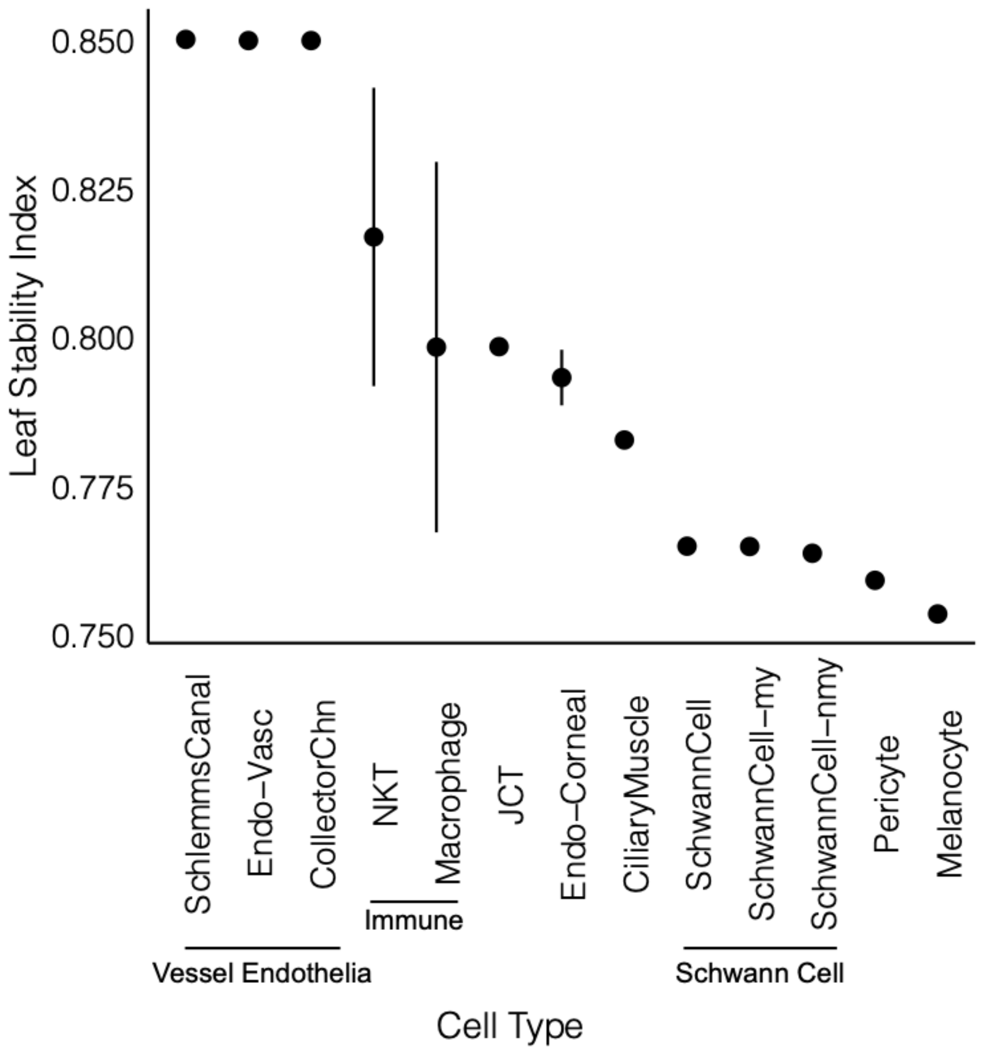


**Figure S3. Leaf stability index by cell type**. Leaf stability indices are highly concordant, with negligible spread within each cell type clade, with the exception of immune cells (NKT and macrophages) and corneal endothelial cells. Mean leaf stability index for each labeled cell type is plotted, vertical lines indicate standard deviation. Superclades are indicated with horizontal lines. Endo-Vasc, vascular endothelium, CollectorChn, collector channel cell, NKT, natural killer T cell, JCT, juxtacanalicular tissue, Endo-Corneal, corneal endothelium, my, myelinating, nmy, non-myelinating.

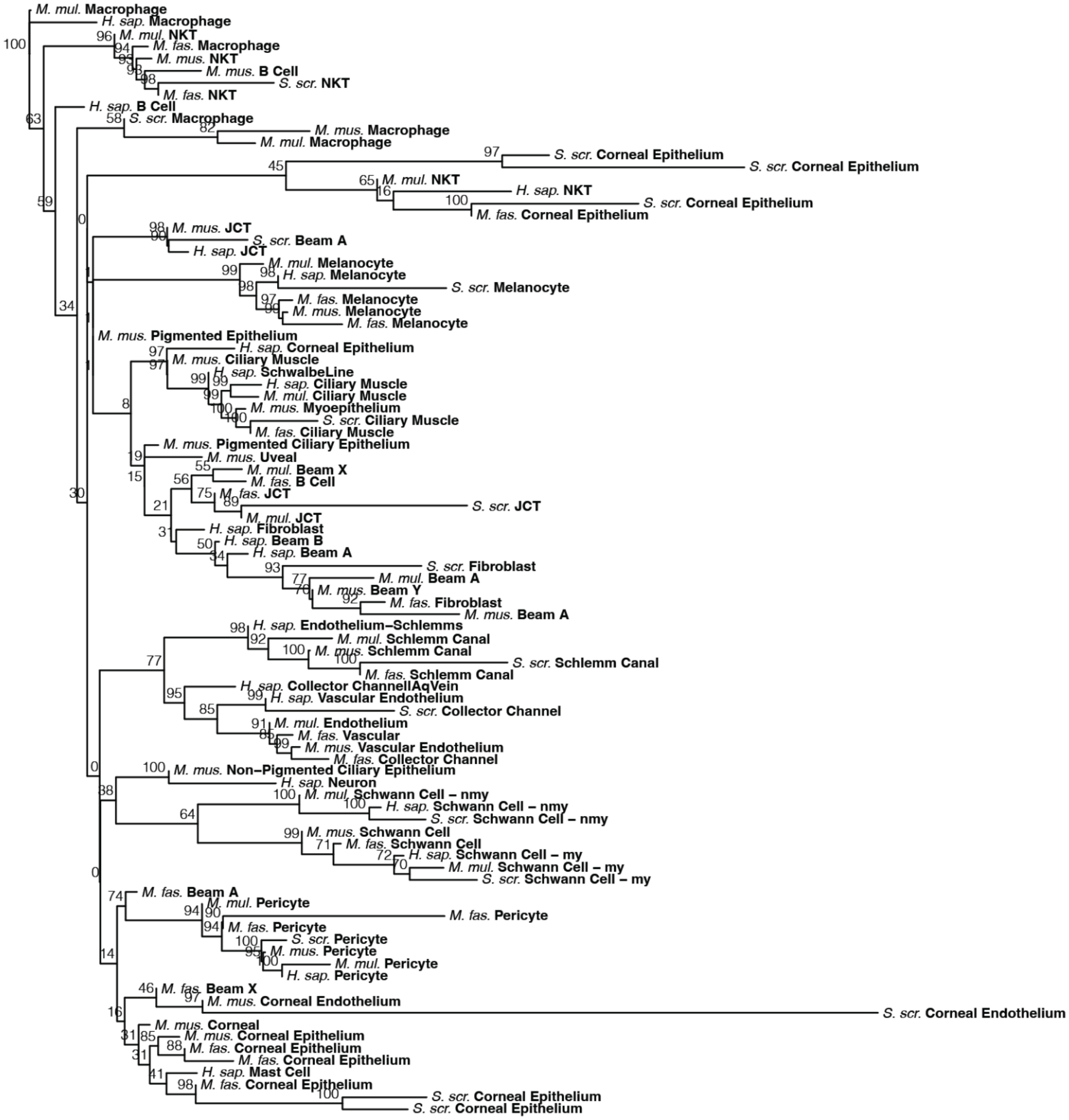


**Figure S4. A cell phylogeny of 92 aqueous humor cells.** This phylogeny was created using the same methods as the 54 cell phylogeny of Figure 4, but was inferred from the 92 cell 20 PC matrix (Fig. S1, step C1.1) that included one cell per cell type group per species for 92 cell type groups. This matrix is more encompassing than the 54 cell 20 PC matrix (Fig. S1, step C1.2), as it includes the unstable cell type groups that were excluded from the final analysis (Methods). Jumble scores are plotted at the nodes. H. sap., *Homo sapiens*, M. fas., *Macaca fascicularis*, M. mul., *Macaca mulatta*, M. mus., *Mus musculus*, Sus. scr., *Sus scrofa*, my, myelinating, nmy, non-myelinating, JCT, juxtacanalicular tissue, NKT, natural killer T cell.

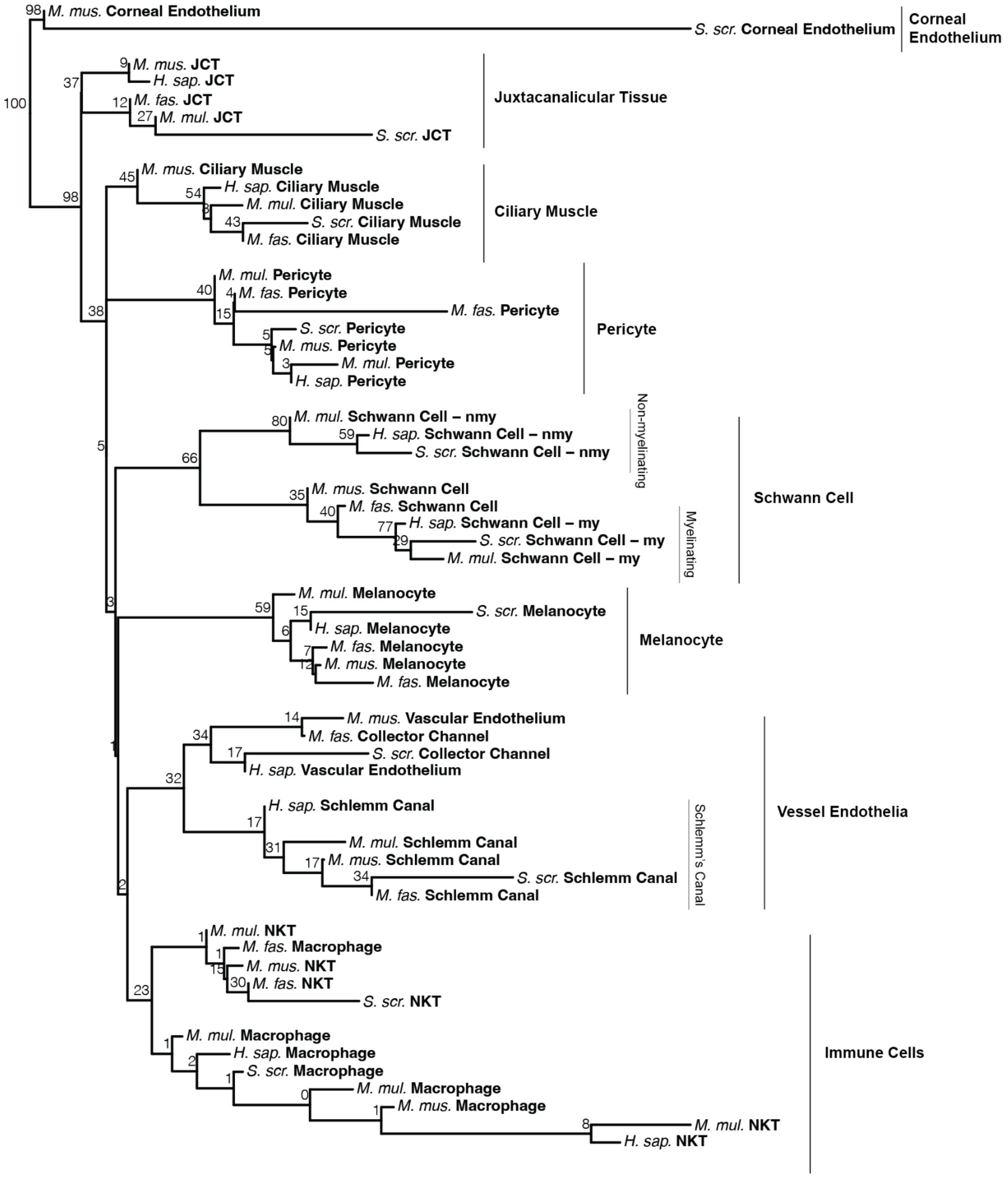


**Figure S5. Felsenstein's bootstrap is not a suitable measure of biological repeatability for a cell phylogeny.** The 54 cell focal phylogeny from Figure 4 is annotated with Felsenstein's bootstrap values calculated from the scjackknife trees. While cell type clades still emerged within any scjackknife tree, cell clades between scjackknife trees often varied by one or a few cells. This subtle degree of variability is not well captured by traditional bootstrap scores, which mark clades only as present or absent based on the presence of all tips, no matter the degree of similarity of the clades.

| PC1 | PC2 | PC3 | PC4 | PC5 | PC6 | PC7 | PC8 | PC9 | PC10 |
|---|---|---|---|---|---|---|---|---|---|
| **FCER1G** | **EMCN** | SPTBN1 | **COL1A2** | CD9 | CRISPLD2 | MGARP | COL4A4 | ANLN | FAM167B |
| **SRGN** | **VWF** | SERPINF1 | **PCOLCE** | **SNCA** | ITIH5 | PERP | SLIT3 | ECT2 | **ADGRL4** |
| **LCP1** | **PTPRB** | COL1A2 | **COL1A1** | **PRNP** | GJA4 | NQO1 | ALCAM | C1QC | **JAM2** |
| **PTPRC** | **RHOJ** | FBLN2 | **DCN** | **ADAM23** | EBF1 | TPT1 | FGF10 | **CD14** | **VEGFC** |
| **CTSS** | **CDH5** | RAMP2 | MGP | **CHL1** | RASD1 | LGALS3 | TMEM178A | **CD163** | **NOS3** |
| **CORO1A** | **PECAM1** | SERPING1 | **APOD** | TSC22D4 | C1QTNF1 | GYG1 | THBS1 | SMC2 | **FLT1** |
| **CD74** | IPO11 | ANXA5 | OGN | TSPAN15 | BTG2 | TKT | CRIM1 | **CSF1R** | **CD34** |
| **RGS1** | **RAMP2** | SEMA3D | **LUM** | **MBP** | BGN | IRF6 | CDH2 | GPR34 | LIFR |
| **TNFAIP3** | **APOLD1** | SULF1 | IGFBP6 | PERP | RGS5 | PHLDA2 | COL4A5 | **MRC1** | **RGS4** |
| **TPT1** | STX11 | OGN | MEDAG | **MEGF9** | GSN | SPINT2 | COL8A2 | CKAP2 | **ENG** |

**Table S1. The most positively loaded genes for early principal components (PCs).** Genes contributing to the cell type-specific signals described in Table 1 are bolded. See Supplementary Files S1 and S3 for descriptions of gene functions and references.

| PC1 | PC2 | PC3 | PC4 | PC5 | PC6 | PC7 | PC8 | PC9 | PC10 |
|---|---|---|---|---|---|---|---|---|---|
| **CALD1** | S100A1 | **MYH11** | **TYR** | DAB2 | NPNT | **SNCA** | DKK2 | CD52 | TFPI |
| **MYL9** | CTSD | **MYOM1** | **TRPM1** | **CTSB** | PDK3 | **PMP22** | ADGRD1 | **CST7** | GPM6A |
| FLNA | **SERPINF1** | **DSTN** | **PMEL** | **CSF1R** | DTNA | TSC22D4 | MFAP5 | CXCR4 | RASGRP3 |
| **TAGLN** | ABHD12 | **CNN1** | TSPAN10 | **MRC1** | TMEM38A | **SEMA3B** | ECM1 | **RUNX3** | SEMA3D |
| **TPM2** | CRABP2 | **ACTA2** | **MLPH** | CTSZ | BDNF | **CNP** | DCN | **LSP1** | ITGA9 |
| **MYLK** | CPXM2 | **CAP2** | **MITF** | **CD83** | TPBG | **ADAM23** | SEMA3E | LIMD2 | TSPAN18 |
| SYNPO2 | ABCA9 | **KCNMB1** | **EDNRB** | C1QC | PLCB4 | **CDH19** | FGL2 | **CD48** | RGS16 |
| **MYH11** | **NDNF** | **PPP1R12B** | **GPNMB** | **CD163** | P3H1 | **MBP** | MEDAG | **AOAH** | LYSMD2 |
| **ACTA2** | **PLP1** | **MYLK** | CTSD | **LGMN** | F5 | IQGAP2 | SEMA3C | ITGA4 | GALNT1 |
| **CSRP1** | **SDC2** | **ATP2A2** | **FMN1** | **FCER1G** | AGL | TXNIP | PCOLCE2 | **TRAF3IP3** | TMTC1 |

**Table S2. The most negatively loaded genes for early principal components (PCs).** Genes contributing to the cell type-specific signals described in Table 1 are bolded. See Supplementary Files S2 and S3 for descriptions of gene functions and references.
